# Genetic Deletion of ASIC3 Alters Left Ventricular Remodeling and Autonomic Function After Myocardial Infarction in Mice

**DOI:** 10.1101/2025.09.03.674120

**Authors:** Karley M. Monaghan, David D. Gibbons, Chad C. Ward, Maram El-Geneidy, William J. Kutschke, Kathy A. Zimmerman, Donald A. Morgan, Harald M. Stauss, Anne Marie S. Harding, Michelle C. M. Bader, Peter M. Snyder, Rasna Sabharwal, Robert M. Weiss, Kamal Rahmouni, Christopher J. Benson

## Abstract

Chronic overactivation of neurohormonal systems is the principal driver of adverse cardiac remodeling following myocardial infarction (MI). Recent data suggest that ablating cardiac afferent neurons in rats attenuates left ventricular (LV) remodeling following MI by blocking this overactivation. Our lab has shown that acid-sensing ion channels (ASICs) are highly expressed in cardiac afferents and may sense myocardial acidosis. We hypothesized that genetic deletion of ASICs might abrogate disadvantageous remodeling after MI by disrupting afferent signaling pathways otherwise resulting in overactivation of neurohormonal responses. To test this hypothesis, we induced MI by coronary artery ligation in wild type (WT) and ASIC3^-/-^ mice and assessed cardiac remodeling by serial echocardiography. We found that ASIC3^-/-^ mice had less LV dilation relative to MI size, increased LV mass, and increased stroke volume compared to WT mice after MI. To investigate a potential role of the autonomic nervous system, we measured renal and splanchnic sympathetic nerve activity, heart rate and systolic blood pressure variability (sBPV), and hemodynamic responses to atropine and propranolol. In addition, we assessed baroreceptor-heart rate and baroreceptor-renal sympathetic nerve activity (RSNA) reflex function. Following MI, ASIC3^-/-^ mice had lower baroreceptor-RSNA reflex sensitivity than WT mice, associated with elevated sBPV. Importantly, sBPV correlated significantly with post-MI changes in LV mass in ASIC3^-/-^ but not WT mice. Our data shows that ASIC3 plays an important role in cardiac remodeling after MI potentially via modulation of baroreflex sensitivity and sBPV. ASIC3 may be further investigated as a potential therapeutic target in heart failure.

## INTRODUCTION

Myocardial infarction (MI) most commonly occurs from sudden coronary artery occlusion and blockage of blood flow. If blood flow is not rapidly restored, the downstream myocardium undergoes both apoptotic and necrotic myocardial cell death (1, 2). Therapeutic advancements, including early restoration of coronary blood flow, have led to remarkably improved outcomes after MI. This has led to more surviving patients who, unfortunately, can go on to develop progressive left ventricular (LV) dysfunction and heart failure. In fact, ischemic heart disease is the leading cause of heart failure (3).

The development of heart failure after MI is determined by the initial infarct size and subsequent cardiac remodeling. Cardiac remodeling is the process of structural and functional changes that lead to a progressively dilated LV and impaired contractile function. Early phase ventricular remodeling occurs within the first few days after infarction and involves regional thinning and expansion of the infarct zone, and ventricular dilation (4, 5). This in turn causes an increase in wall stress of surviving myocardium in both the border zone surrounding the infarct and remote areas. This elevated mechanical stress, together with local paracrine/autocrine factors and activation of systemic neurohormonal reflexes, can lead to late phase remodeling characterized by hypertrophy of surviving myocytes and further ventricular chamber dilation (6–8). While myocyte hypertrophy can be an adaptive response to offset wall stress and maintain stroke volume, excessive LV remodeling and dilation is a major determinant of whether patients will go on to manifest heart failure, and is one of the strongest predictors of mortality after MI (9, 10).

The decrease in cardiac function and local mechanical/biochemical changes within the myocardium after MI trigger the activation of neurohormonal systems including the sympathetic nervous system, the renin-angiotensin aldosterone systems, as well as release of natruiretic peptides from the heart. In the short term, this can help to maintain cardiac output and perfusion of critical organs after MI. However, chronic activation of these neurohormonal systems has been shown to cause adverse ventricular remodeling and the progression to heart failure (11). Over the past 50 years, tremendous progress has been made in understanding the detrimental role that these neurohormonal changes play in triggering ventricular remodeling. Therapies targeting these systems have become the cornerstone of treatment after MI – resulting in a reduction of LV volume and mass, increased ejection fraction, and decreased mortality (12, 13). While the efficacy of neurohormonal modulation is proven, few new therapies have come forth within the past couple of decades.

Recent studies in animal models of heart failure suggest a new approach. Rather than targeting the efferent limbs of these reflexes, inhibition of the sensory (afferent) pathways that trigger these neurohormonal pathways have shown beneficial effects on cardiac remodeling and cardiovascular function (14, 15). Most of this work has focused on the sensory neurons that innervate the heart, and particularly those cardiac afferents that follow the sympathetic nerve pathways back to their cell bodies located in the dorsal root ganglia (termed ‘cardiac sympathetic afferents’). Besides signaling pain during myocardial ischemia or infarction, cardiac sympathetic afferents also trigger sympathoexcitation (16, 17). This ‘cardiac sympathetic afferent reflex’ is enhanced in heart failure and is a major contributor to sympathoexcitation in heart failure (18–20). In a rat model of heart failure after MI, the group of researchers around Dr. Irvine H. Zucker found that intrapericardial injection of resiniferotoxin to ablate (denervate) cardiac sympathetic afferents prevented sympathoexcitation, improved baroreflex sensitivity, and also reduced cardiac dilation and improved cardiac diastolic function (14, 21). However, the molecular receptors within cardiac afferents that are activated in the setting of myocardial ischemia and heart failure are incompletely understood. Interestingly, the ion channel TRPV1, which is the target of the toxin resiniferotoxin used to ablate cardiac afferents in the above mentioned study, does not appear to be the sensor since TRPV1 null mice actually show worsened inflammation, fibrosis, and deleterious remodeling after MI (22, 23).

As an organ with high metabolic activity, the heart is susceptible to rapid drops in pH during myocardial ischemia (24), and acidosis is a potent activator of cardiac sympathetic afferent fibers in vitro (25, 26). In isolated labeled cardiac sympathetic afferent neurons we found that acid solution evoked much larger currents and were present in a larger percentage of cardiac afferents than other potential chemical agonists, and their current properties were consistent with acid-sensing ion channels (ASICs) (27, 28). ASICs belong to the DEG/ENaC family of ion channels that include four genes (*ASIC1, -2, -3,* and *-4*) encoding 6 subunits (ASIC1a, -1b, - 2a, -2b, -3, -4) in rodents. Functional ASIC channels consist of a complex of three subunits that can form homomultimeric channels with three of the same subunits or heterotrimers composed of two or more subunits (29, 30). Besides being activated by interstitial acidosis, ASICs are activated or potentiated by other metabolites that are released by ischemic tissue including lactate, arachidonic acid, and ATP – suggesting that ASICs may serve as metabolic and pain sensors during stress conditions such as ischemia (31–33). We have subsequently shown that the composition of ASIC channels in murine cardiac sympathetic afferent neurons is heteromeric composed of ASIC2a and ASIC3 subunits, with ASIC3 being responsible for its exquisite pH sensitivity in the ranges that occur during myocardial ischemia (34, 35).

Given that 1) cardiac afferents are important triggers of sympathoexcitation and deleterious cardiac remodeling after MI, 2) ASIC3 is highly expressed and is a sensitive pH sensor within cardiac afferents, and 3) ASIC3 is generally coexpressed in the same set of sensory neurons that express TRPV1 (the target of resiniferotoxin) (36), we tested if genetic deletion of ASIC3 would alter cardiac remodeling and function after MI.

## METHODS

### Ethical Approval

All experimental procedures and protocols were approved by the Institutional Animal Care and Use Committee of the University of Iowa (Protocol no. 2051770) and conformed to the national guidelines set by the Association for Assessment and Accreditation of the Laboratory Animal Care.

### Animals

ASIC3^-/-^ mice were generated (37) and bred on a C57Bl/6J background and were compared to wild-type (WT) C57Bl/6J mice (Jackson Laboratories). These mice were subsequently backcrossed for 10 generations onto a C57Bl/6J background to generate a congenic line.

Following the initial development of the strain, the ASIC3^-/-^ mouse line was backcrossed every 10 generations to avoid genetic drift. All mice were housed at the University of Iowa in a temperature-controlled room (22C°) with a 12-hour light/dark cycle with ad libitum access to both water and standard mouse chow. Both male and female mice were used in all groups.

Since there were no sex differences in cardiac remodeling after MI, we analyzed them together (Supplemental Table 1). Mice were euthanized with CO_2_ per AVMA guidelines.

### Echocardiography

Cardiac function was measured at baseline by echocardiography in 7-week-old, lightly-sedated mice (midazolam 0.1 mg SQ injection) using a Vevo 2100 instrument (Visualsonics, Toronto, Canada) equipped with a 30-MHz probe as previously described (38). Briefly, the anterior chest hair was shaved and warmed gel was applied. The mouse was scruffed and held in the left lateral position. Images of the short and long axis were obtained with a frame rate of ∼180-250 Hz. All image analysis was performed using Vevo 2100 analysis software version 5.7.1, by an investigator blinded to genotype. Endocardial and epicardial borders were traced on the short axis view at end-diastole and end-systole. LV length was measured from endocardial and epicardial borders to the LV outflow tract in end-diastole and end-systole. The biplane area-length method was performed to calculate LV volumes and mass. We previously found a tight correlation between echocardiographic and isolated and weighed LV mass (39). 48 hours after surgery echocardiograms were performed without sedation to evaluate the ligation-induced myocardial injury. IZ fraction was reported as the non-contractile fraction of the LV. Three weeks post-surgery another echocardiogram was done without sedation to evaluate cardiac remodeling. Previous data in mice after coronary ligation show that most of the cardiac remodeling occurs by 2.5 weeks and doesn’t change much through 6 weeks post-MI (40).

### MI/Sham Surgery

One week following baseline echocardiograms, ASIC3^-/-^ and WT mice (8 weeks of age) underwent MI or sham surgery as previously described (41). Animals were anesthetized with isoflurane, prepared, and ventilated. Hearts were accessed via a thoracotomy and infarcted by placing a permanent ligature on the mid-left anterior descending coronary artery using aseptic technique. The ribcages were then closed with the air evacuated, and the incision closed. The sham procedure followed the same protocol while omitting the artery ligation. Mice were then monitored continuously until awake, and analgesia was administered immediately and daily for 2 days postoperatively. Mice were returned to their home cages with sex-matched litter mates and monitored two times daily for 5 days. Mice that underwent MI surgery but did not develop an MI according to echocardiography were excluded.

### Radiotelemetry recording of hemodynamics

One week after MI/Sham surgery, radio telemeters were implanted in a cohort of mice as previously described (42). Briefly, under isoflurane anesthesia a pressure-sensing implantable mouse transmitter (TA11PA-C10, DSI) was inserted in the left common carotid artery and advanced into the aorta through a midline incision. The body of the transmitter was tunneled subcutaneously to the left flank, and the incision closed. Following surgery, the analgesic flunixin meglumine (0.05 µg/g) was injected subcutaneously. The mice were individually housed and monitored daily. After the 3-week echocardiogram, cages were transferred to the telemeter room and allowed to acclimate for two days before turning on telemeters. Baseline heart rate and blood pressure were continuously recorded for 48 hours. Following the baseline recordings, cardiovagal and cardiac sympathetic tone were measured from HR responses to methylatropine (1 mg/kg, intraperitoneal [i.p.]) and propranolol (1 mg/kg, i.p.) respectively.

Autonomic modulation of cardiac function was also assessed via heart rate variability using both the root mean square of successive differences (RMSSD, a measure of vagal modulation) and power spectral analysis from the baseline recordings before drug injections (43). For spectral analysis, the low frequency (LF) band was set at 0.2-0.8 Hz and the high frequency (HF) band was set at 0.8-5.0 Hz. Baroreceptor-heart rate reflex sensitivity was calculated using the sequence technique as previously described (44). Briefly, sequences of three or more consecutive heart beats where both arterial blood pressure and interbeat intervals simultaneously increased or decreased were detected. Linear regressions were calculated for these sequences.

### Sympathetic nerve recording

In a sub cohort of mice following 3-week echocardiograms, regional sympathetic nerve activity (SNA) was measured using direct multifiber recording in anesthetized mice as previously described (45). Mice were anesthetized with i.p. injection of ketamine (91 mg/kg) and xylazine (9.1 mg/kg). Each mouse was intubated (PE-50) to allow for spontaneous respiration of oxygen-enriched room air. Body temperature, measured with a rectal probe, was kept constant at 37.5°C by using a surgical heat lamp and a metal heat platform. The left jugular vein was cannulated with a micro-renathane tubing (MRE-40) to sustain the level of anesthesia throughout the 4-hour protocol with α-chloralose (initial dose: 12 mg/kg; sustaining dose of 6 mg/kg/h) and for drug treatments. Finally, the left carotid artery was cannulated with a tapered micro-renathane tubing (MRE-40) for continuous measurement of arterial pressure and heart rate.

A dissecting microscope was employed: the nerves subserving the left kidney (RSNA) and visceral organs (splanchnic SNA) were identified, carefully dissected free, and placed on a bipolar 36-gauge platinum-iridium electrode (A-M Systems). When the optimum recording of SNA was obtained from each nerve, the electrode was covered with silicone gel (Kwik-Sil; World Precision Instruments Inc). The electrode was attached to a high-impedance probe (HIP-511, Grass Instruments), and the nerve signal was amplified 10^5^ times with a Grass P5 AC pre-amplifier. The amplified nerve signal was filtered at a 100-Hz and 1000-Hz and was then routed to a speaker system and to an oscilloscope (model 54501 A, Hewlett–Packard) to monitor the audio and visual quality of the sympathetic nerve recordings. The amplified, filtered nerve signal was also directed to a MacLab analogue-digital converter (Model 8 S, AD Instruments Castle Hill) containing software (MacLab Chart Pro; Version 7.0) that uses a cursor to analyze the number of spikes/second that exceeds the background noise threshold.

Arterial baroreflex control of renal SNA was tested by intravenous infusion of sodium nitroprusside (at doses of 0.05, 0.1, 0.5, and 1 μg) followed by phenylephrine (at doses of 0.05, 0.1, 0.5, and 1 μg), which decreases and increases arterial pressure, respectively. To ensure that background electrical noise was excluded from sympathetic measurements, post-mortem background activity was subtracted from all SNA.

### Quantitative PCR

Whole hearts were extracted and homogenized in TRIzol™ Reagent (Invitrogen, 15596026). RNA was extracted following the manufacturer’s protocol. RNA was quantified using a Nano drop spectrophotometer. Reverse transcription was done using 1 μg of RNA with the High-capacity cDNA Reverse Transcription Kit (Applied Biosystems™, 4368814). Quantitative PCR was done using the *Power* Sybr™ Green PCR Master Mix (ThermoFisher, 4367659) and read on the Applied Biosystems QuantStudio 7 Flex PCR instrument. Data was analyzed using the 2^-ΔΔCt^ method and the WT Sham group as the calibrator. Primer sequences for the target genes are listed in the table below. Sequences were taken from the Harvard Medical School PrimerBank unless otherwise indicated.

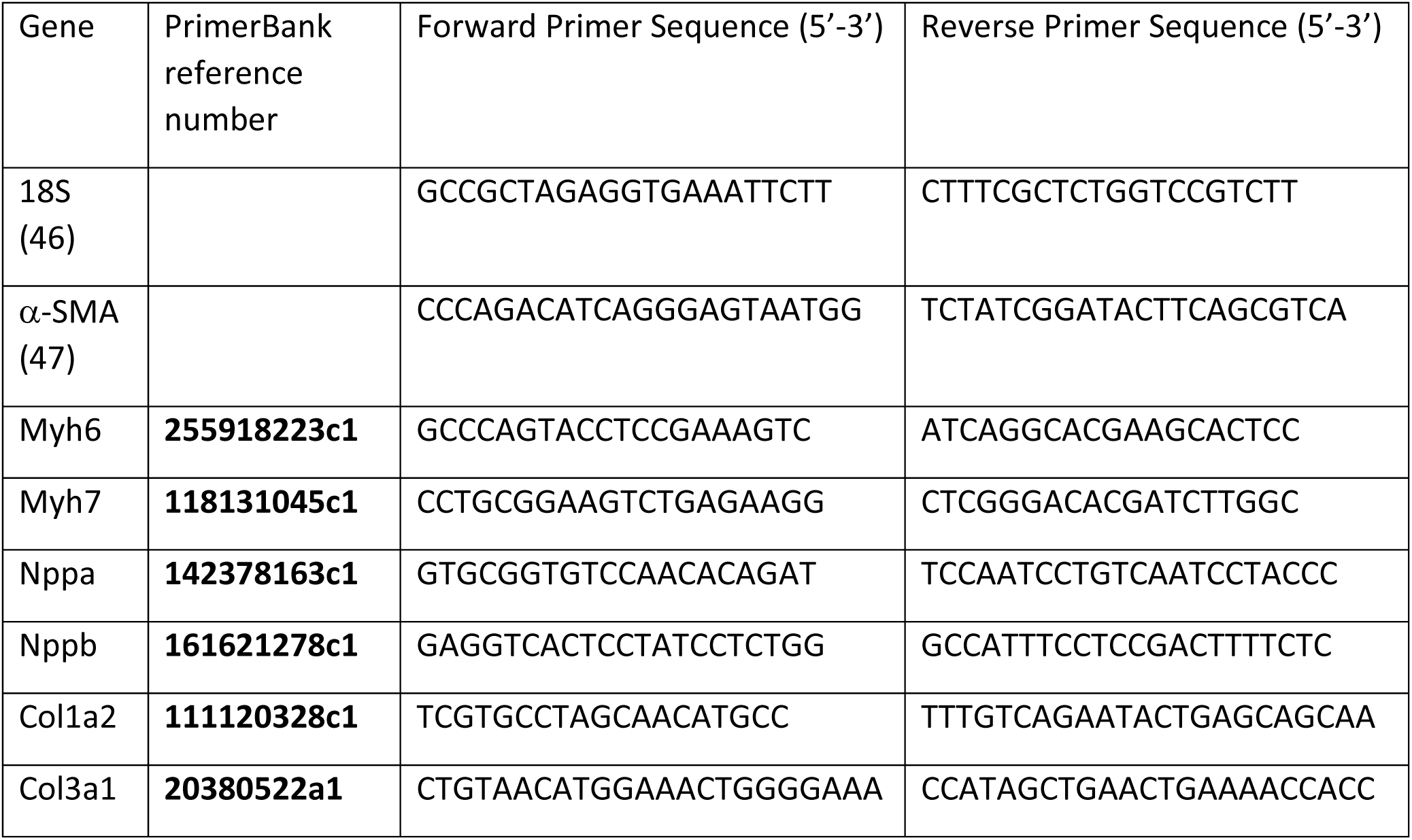

### Statistics

GraphPad Prism (10.4.1) was used to analyze statistical data. Bar graphs represent means ± standard error of the mean (SEM). Three-way ANOVA, two-way ANOVA, unpaired *t*-test, simple linear regression, and chi square were used as appropriate to analyze results. If significant differences were observed, the indicated post hoc tests as described in the figure legends were performed. Significance was reported at *P* < 0.05.

## RESULTS

### ASIC3^-/-^ mice have improved stroke volume and undergo less LV dilation relative to infarct size after MI than WT mice

To test if ASIC3 contributes to cardiac remodeling after MI, baseline echocardiograms were performed prior to MI/Sham surgery and then repeated at 48 hours post-op to confirm the presence and size of infarction (IZ fraction of the LV). Three weeks after surgery a third echocardiogram was performed to quantify cardiac remodeling. Figure 1A shows representative short axis echocardiographic images in systole (*left column*) and m-mode images (*right* column) for a WT Sham (*top*), WT MI (*middle*), and ASIC3^-/-^ MI (*bottom*) heart at 3-weeks post-MI. Compared to baseline, echocardiograms at 48 hours post-MI revealed that both WT and ASIC3^-/-^ MI mice underwent a significant increase in LV end-diastolic volume (LVEDV) (Fig. 1B: *left*) and LV end-systolic volume (LVESV) (Fig. 1C: *left*), a significant decrease in LV ejection fraction (EF) (Fig 1D: *left*), and no change in stroke volume (SV) (Fig. 1E: *left*). While early remodeling can occur within the first 48 hours after MI, most late remodeling happens between 48 hours and 3 weeks after MI in mice (40). Both WT MI and ASIC3^-/-^ MI mice underwent further increases in LVEDV at 3 weeks compared to 48 hours (Fig. 1B: *left*). WT MI mice also had a further significant increase in LVESV between 48 hours and 3 weeks, however the change in LVESV of ASIC3^-/-^ MI mice were not significant (Fig. 1C: *left*). Neither the WT nor the ASIC3^-/-^ MI mice saw changes in EF between 48 hours and 3 weeks (Fig D: *left*). In addition, ASIC3^-/-^ MI mice had a significant increase in stroke volume between 48 hours and 3 weeks, whereas WT mice did not (Fig. 1E: *left*). To further quantify LV remodeling, we compared the change in each parameter between 48 hours and 3 weeks after MI between MI groups. Whereas there were no significant differences between WT and ASIC3^-/-^ mice in ΔLVEDV, ΔLVESV, and ΔLVEF (Figs. 1B, C, and D: *right*), ASIC3^-/-^ MI mice increased their LV stroke volume between 48 hours and 3 weeks more than the WT MI mice (Fig. 1E; *right*).

**Figure 1.**
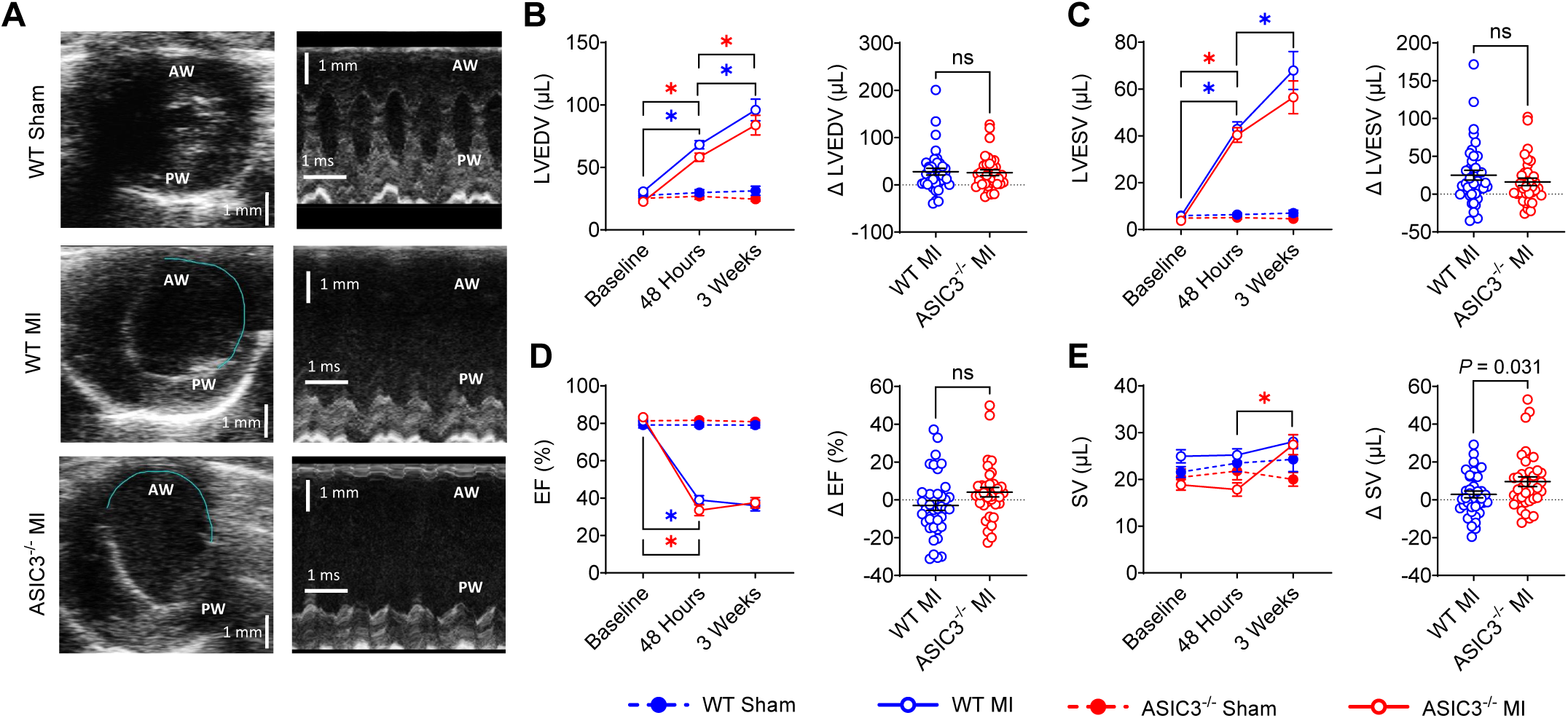
Echocardiographic data. **A)** <u>Left column:</u> Short-axis echocardiograms at end-systole. Teal: endocardial silhouette of akinetic myocardium. Ischemic Zone = path length of akinetic endocardial silhouette/pathlength of total endocardial silhouette. <u>Right column:</u> M-mode echocardiograms. Anterior left ventricular wall (AW), posterior left ventricular wall (PW). **B)** Left Ventricular End Diastolic Volume (LVEDV): Three-way ANOVA revealed a significant effect of time (*F*(_2,230_) = 59.9, *P* ≤ 0.0001), surgery (*F*(_1,115_) = 82.63, *P* ≤ 0.0001) and a significant time x surgery interaction (*F*(_1,115_) = 54.02, *P* ≤ 0.0001). Tukey post-hoc test found significant differences for WT MI between baseline and 48 hours (*P* ≤ 0.0001) and 48 hours and 3 weeks (*P* ≤ 0.0001) and for ASIC3^-/-^ MI between baseline and 48 hours (*P* ≤ 0.0001) and 48 hours and 3 weeks (*P* = 0.0002). **C)** Left Ventricular End Systolic Volume (LVESV): Three-way ANOVA revealed a significant effect of time (*F*(_2,230_) = 60.59, *P* ≤ 0.0001), surgery (*F*(_1,115_) = 97.73, *P* ≤ 0.0001) and a significant time x surgery interaction (*F*(_1,115_) = 58.87, *P* ≤ 0.0001). Tukey post-hoc test found significant differences for WT MI between baseline and 48 hours (*P* ≤ 0.0001) and 48 hours and 3 weeks (*P* = 0.02) and for ASIC3^-/-^ MI between baseline and 48 hours (*P* ≤ 0.0001). **D)** Ejection Fraction (EF): Three-way ANOVA revealed a significant effect of time (*F*(_2,230_) = 201.5, *P* ≤ 0.0001), surgery (*F*(_1,115_) = 345.5, *P* ≤ 0.0001) and a significant time x surgery interaction (*F*(_1,115_) = 200.9, *P* ≤ 0.0001). Tukey post-hoc test found significant differences for WT MI between baseline and 48 hours (*P* ≤ 0.0001) and for ASIC3^-/-^ MI between baseline and 48 hours (*P* ≤ 0.0001). **E)** Stroke Volume (SV): Three-way ANOVA revealed a significant effect of time (*F*(_2,230_) = 6.01, *P* = 0.003), genotype (*F*(_1,115_) = 9.29, *P* = 0.003) and a significant time x surgery interaction (*F*(_1,115_) = 5.14, *P* = 0.007). Tukey post-hoc test found significant differences for ASIC3^-/-^ MI between 48 hours and 3 weeks (*P* = 0.028). Unpaired Student’s *t*-tests revealed a significant difference between MI groups for SV (*P* = 0.031). Values are reported as mean ± SEM. (Nonsignificant (ns), significant difference for WT MI (*), significant difference for ASIC3^-/-^ MI (*); WT Sham: *N* = 21, WT MI: *N* = 39, ASIC3^-/-^ Sham: *N* = 24, ASIC3^-/-^ MI: *N* = 35).

Left anterior descending artery ligation produces a high variability of infarctions in mice, which can lead to a high variability in the degree of remodeling. To investigate the effect of MI size on the cardiac remodeling, we correlated the 3-week echo data to infarct size (see Fig. 1A). As expected, both the WT and ASIC3^-/-^ mice saw a positive correlation between LV volumes at 3 weeks post-MI and infarction size (IZ fraction). When fitted with a simple linear regression, the y-intercepts of the LVEDV (Fig. 2A) and LVESV (Fig. 2C) for the ASIC3^-/-^ mice were significantly lower than for the WT mice. Thus, for any given infarct size, ASIC3^-/-^ mice had less LV dilation.

**Figure 2.**
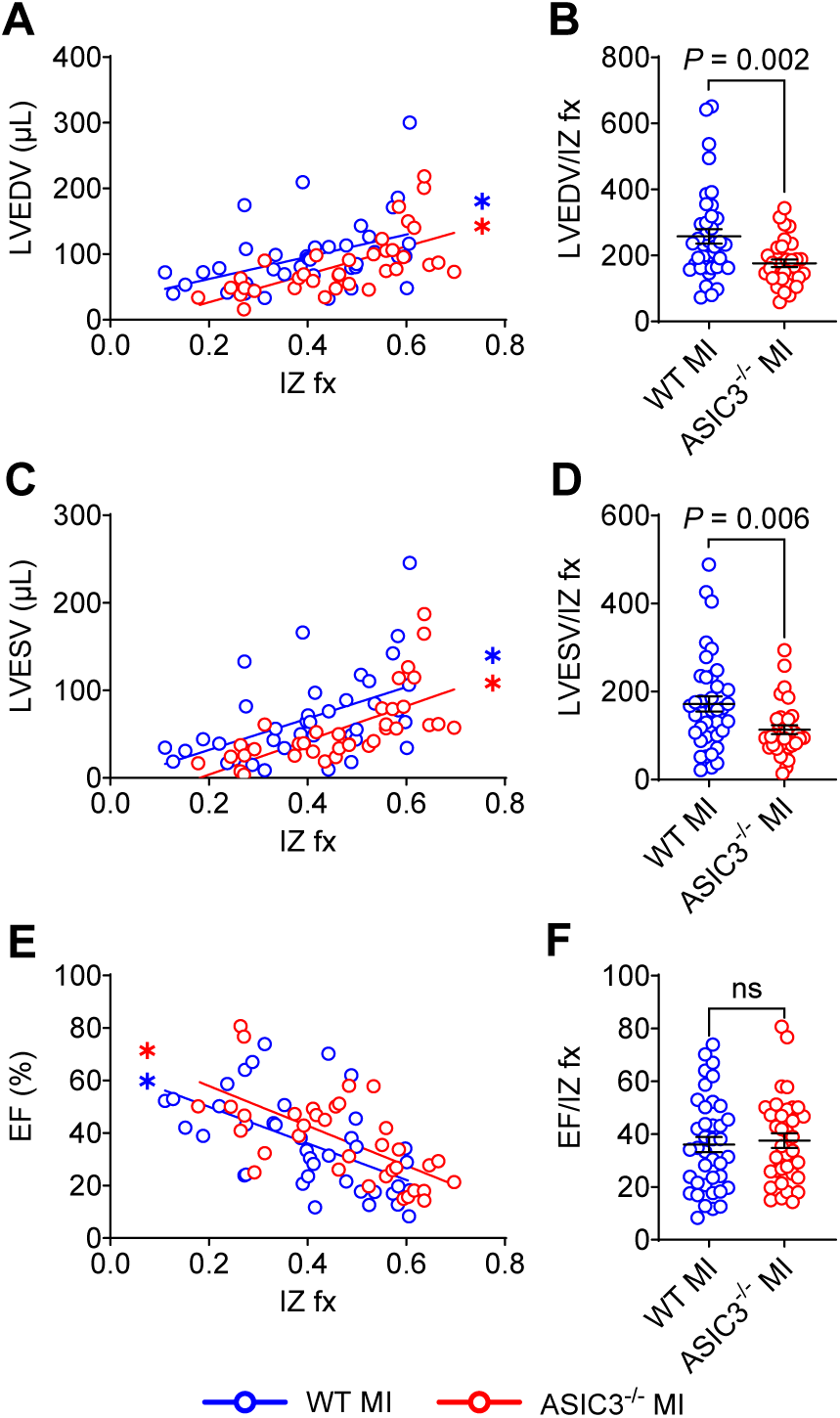
Effect of myocardial infarction (MI) size on cardiac remodeling after MI. **A)** Correlation of left ventricular end diastolic volume (LVEDV) to IZ fx. Simple linear regression testing revealed significant correlations for both WT MI (*R* = 0.44, *P* = 0.0046) and ASIC3^-/-^ MI (*R* = 0.65, *P* ≤ 0.0001), as well as a significant difference in the y-intercepts of the groups (*F*(_1,71_) = 5.89, *P* = 0.018); volume (LVESV) to IZ fx. Simple linear regression testing revealed significant correlations for both WT MI (*R* = 0.5, *P* = 0.001) and ASIC3^-/-^ MI (*R* = 0.67, *P* < 0.0001), as well as a significant difference in the y-intercepts of the groups (*F*(_1,71_) = 6.96, *P* = 0.01). **D)** LVESV normalized to IZ fx. Unpaired *t*-test revealed a significant difference between groups (*P* = 0.006). **E)** Correlation of left ventricular ejection fraction (LVEF) to IZ fx. Simple Linear Regression testing revealed significant correlations for both WT MI (*R* = 0.56, *P* = 0.0002) and ASIC3^-/-^ MI (*R* = 0.65, *P* ≤ 0.0001), as well as a trend towards a significant difference in the y-intercepts of the groups (*F*(_1,71_) = 3.82, *P* = 0.055). **F)** LVEF normalized to IZ fx. Values are reported as Mean ± SEM. (Nonsignificant (ns), significant correlation for WT MI (*), significant correlation for ASIC3^-/-^ MI (*); WT MI: *N* = 39, ASIC3^-/-^ MI: *N* = 35).

Correlation linear regression fits of LV EF to infarction size show that ASIC3^-/-^ MI mice were not statistically different from the WT MI mice but trended (*P* = 0.055) towards greater LV EF compared to WT mice after MI (Fig. 2E). Additionally, we normalized the 3-week echocardiogram data with the IZ fraction (Figs. 2B, D, and F) and found LVEDV and LVESV to be lower in ASIC3^-/-^ mice compared to WT mice. In summary, these results indicate that ASIC3^-/-^ mice undergo less LV dilation and have improved stroke volume compared to WT mice after MI.

### LV mass increases in ASIC3^-/-^ but not WT mice after MI

As a measure of hypertrophy, we measured LV mass via echocardiography at baseline and at 48 hours and 3 weeks after MI. Between 48 hours and 3 weeks, ASIC3^-/-^ MI mice increased their LV mass significantly more than WT MI mice and were the only group to undergo an increase in mass between 48 hours and 3 weeks (Fig. 3A). Simple linear regression of the correlation between LV mass at 3 weeks post-MI and IZ fraction showed a positive correlation in the ASIC3^-/-^ MI but not in WT MI mice (Fig 3B). This implies that ASIC3^-/-^ MI mice underwent increasing LV hypertrophy per increase in MI size, whereas this did not occur in WT MI mice. To validate that the increase in mass wasn’t due to differences in animal size we normalized these results to body weight in a subset of animals for which we had body weights at all three time points (Fig. 3C and Supplemental Table 2). ASIC3^-/-^ MI mice had a significant increase in LV mass when normalized to body weight between baseline and 3 weeks, whereas no other groups saw an increase (Fig. 3C). When normalized to body weight, ASIC3^-/-^ MI mice had greater LV mass at 3 weeks compared to all other groups (Fig. 3D), and the change in normalized LV mass at 3 weeks compared to 48 hours was also greater in ASIC3^-/-^ mice than in WT (Fig. 3E).

**Figure 3.**
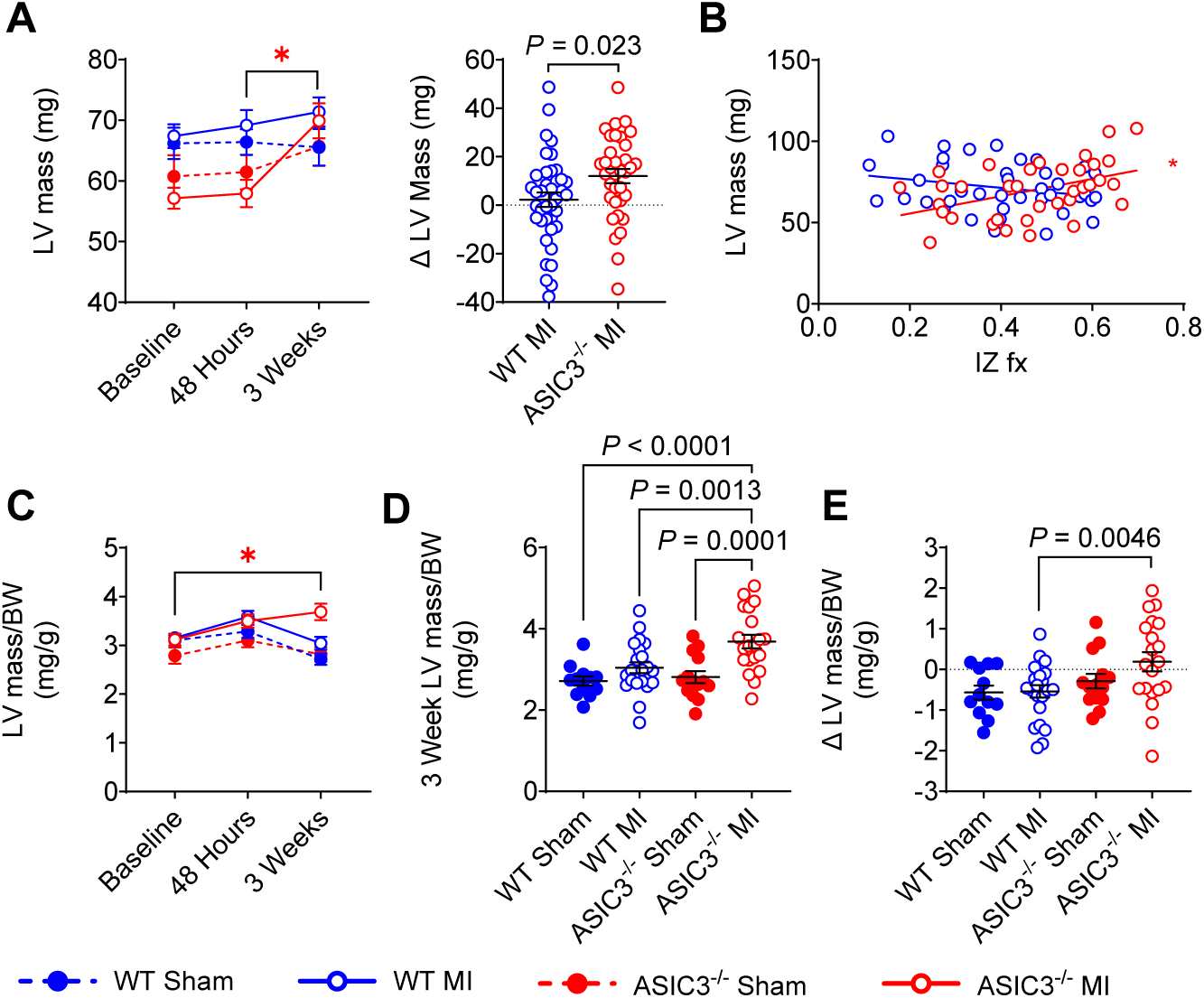
Change in left ventricular (LV) mass after myocardial infarction (MI). **A)** Left: LV mass measured from echocardiograms plotted over time in WT (blue) and ASIC3^-/-^ (red) mice after sham (closed circles) or MI (open circles) surgery. Three-way ANOVA revealed a significant effect of time (*F*(_2,230_) = 6.93, *P* = 0.001), genotype (*F*(_1,115_) = 7.51, *P* = 0.007) and a significant time x genotype interaction (*F*(_2,230_) = 3.88, *P* = 0.022). Tukey post-hoc test found significant differences for ASIC3^-/-^ MI between 48 hours and 3 weeks (*P* = 0.011). Right: the change in LV mass from 48 hours post-MI to 3 weeks post-MI in WT and ASIC3^-/-^ mice after sham or MI surgery. Unpaired *t*-test revealed significant differences between MI groups (*P* = 0.023, *). **B)** Correlation between 3-week LV mass and ischemic zone fraction (IZ fx) in WT and ASIC3^-/-^ mice. Simple Linear Regression testing revealed a significant correlation for ASIC3^-/-^ MI (*R* = 0.442, *P* = 0.0078, *) but not WT MI (*R* = 0.24, *P* = 0.14), and a significant difference in the slopes of the groups (*F*(_1,70_) = 9.84, *P* = 0.0025); lines are best fit regressions (WT Sham: *N* = 21; WT MI: *N* = 39, ASIC3^-/-^ Sham: *N* = 24; ASIC3^-/-^ MI: *N* = 35). **C)** LV mass normalized to body weight (BW) at indicated intervals in a subset of mice in which body weight was obtained at each time point. Three-way ANOVA revealed a significant effect of time (*F*(_2,128_) = 7.46, *P* = 0.0009), surgery (*F*(_1,64_) = 17.42, *P* ≤ 0.0001) and a significant time x genotype interaction (*F*(_2,128_) = 5.15, *P* = 0.007). **D)** LV mass at 3 weeks normalized to body weight for data in **C**. Two-way ANOVA revealed a significant effect of surgery (*F*(_1,64_) = 14.9, *P* ≤ 0.0001) and genotype (*F*(_1,64_) = 5.69, *P* = 0.020). Fisher’s LSD post-hoc test found significant differences between ASIC3^-/-^ MI and WT Sham (*P* ≤ 0.0001), WT MI (*P* = 0.0013), and ASIC3^-/-^ Sham (*P* = 0.0001). **E)** Change in LV Mass normalized to BW between 48 hours and 3 weeks post-MI for data in **C**. Two-way ANOVA revealed a significant effect of genotype (*F*(_1,64_) = 6.31, *P* = 0.015). Fisher’s LSD post-hoc test found a significant difference between ASIC3^-/-^ MI and WT MI (*P* = 0.0046). Values are reported as mean ± SEM. (WT Sham: *N* = 12; WT MI: *N* = 22, ASIC3^-/-^ Sham: *N* = 14, ASIC3^-/-^ MI: *N* = 20).

### Altered neurohormonal responses in ASIC3^-/-^ mice compared to WT mice after MI

To understand the underlying mechanisms that lead to altered cardiac remodeling in ASIC3^-/-^ mice compared to WT mice after MI, we first recorded hemodynamics continuously for two days in conscious mice in their home cage using the radiotelemetry system. The average heart rate over the two days did not differ between any of the groups (Fig. 4A). The mean arterial blood pressure (MAP) in the WT MI mice was significantly lower than the WT Sham group. In contrast, there were no differences in MAP between the ASIC3^-/-^ Sham and ASIC3^-/-^ MI. While there was no significant difference in BP between the MI groups over the 48-hour period (Fig. 4B), ASIC3^-/-^ MI mice had higher MAP and systolic blood pressure (BP) at night than WT MI mice (Supplemental Table 3).

**Figure 4.**
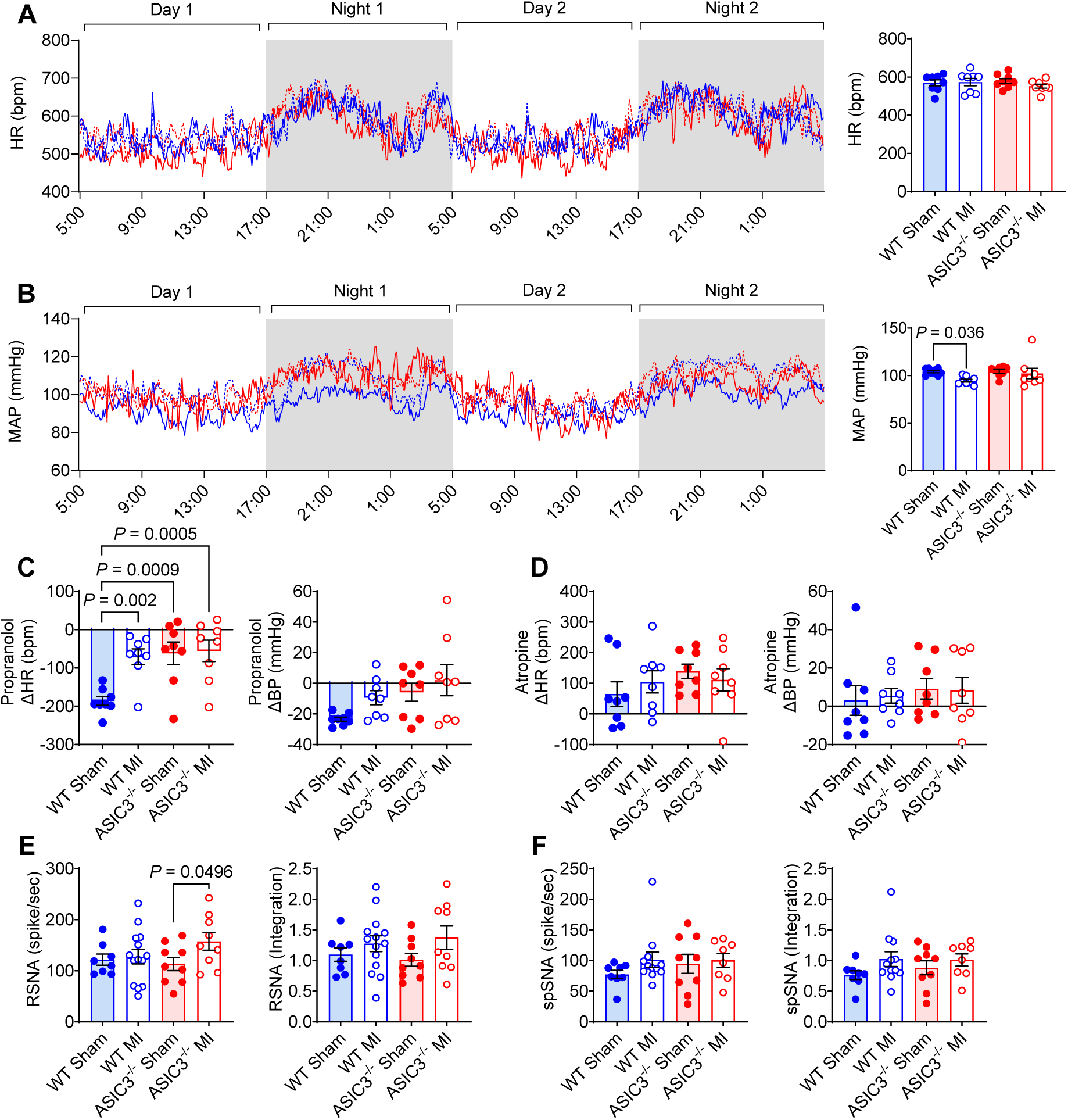
Measure of autonomic nervous system tone. **A)** Average heart rate of radio-telemeter implanted wild-type (WT; blue) and ASIC3^-/-^ (red) mice over 2 days. Traces of heart rate over the 2-day period (*left*) and average of the two days (*right*). Two-way ANOVA of HR data with Fisher’s LSD post-hoc adjustment showed no significant effects. **B)** Mean arterial pressure (MAP) of radio-telemeter implanted WT and ASIC3^-/-^ mice over 2 days. Traces of BP over the 2-day period (*left*) and average of the two days (*right*). Two-way ANOVA with Fisher’s LSD post-hoc adjustment of MAP data showed no significant effects.A significant difference was observed between WT Sham and WT MI. **C)** Change in heart rate (HR, *left*) and blood pressure (BP, *right*) after injection of 0.1 mg/kg propranolol. Two-way ANOVA revealed a significant effect of surgery (*F*(_1,28_) = 4.68, *P* = 0.039). Fisher’s LSD post-hoc test found significant differences between WT Sham and WT MI, WT Sham and ASIC3^-/-^ Sham, and WT Sham and ASIC3^-/-^ MI. **D)** Change in HR (*left*) and BP (*right*) after injection of 0.1 mg/kg atropine. Two-way ANOVA with Fisher’s LSD post-hoc test found no significant effects or individual differences. (*N* = 8 for each group). **E)** Renal sympathetic nerve activity (RSNA). Two-way ANOVA with a Fisher’s LSD post-hoc test found a significant difference between ASIC3^-/-^ Sham and ASIC3^-/-^ MI. (WT Sham: *N* = 8; WT MI: *N* = 14, ASIC3^-/-^ Sham: *N* = 9, ASIC3^-/-^ MI: *N* = 9). **F)** Splanchnic sympathetic nerve activity (spSNA). Two-way ANOVA with Fisher’s LSD post-hoc test found no significant effects or individual differences. (WT Sham: *N* = 8; WT MI: *N* = 12, ASIC3^-/-^ Sham: *N* = 9, ASIC3^-/-^ MI: *N* = 8).

Next, we evaluated baseline parasympathetic and sympathetic autonomic tone. As a measure of cardiac sympathetic tone in conscious mice we recorded HR and BP responses, respectively, to acute injection of propranolol (β-adrenergic receptor blocker). Propranolol decreased HR in all groups, with a significantly larger decrease in the WT sham mice compared to all other groups (Fig. 4C, *left*). BP was also decreased in the WT sham mice, but no differences were seen between the groups (Fig. 4C, *right*). As a measure of cardiac vagal tone, acute injection of atropine (muscarinic cholinergic receptor blocker) increased HR in all groups and had a marginal effect on BP, but no differences were seen between the groups following atropine injection (Fig. 4D). In a separate cohort of anesthetized mice, we directly recorded sympathetic nerve activity (SNA) from both the renal and splanchnic nerves. ASIC3^-/-^ mice displayed a slight elevation in the renal SNA spike frequency following MI (Fig 4E, *left*) with no differences in the overall (mean rectified/integrated) activity (Fig. 4E, *right*), and the MI groups did not differ. Unexpectedly we did not see a significant increase in renal SNA in the WT mice following MI. There were also no differences between the groups in splanchnic SNA (Fig. 4F).

As additional measures of autonomic dysregulation, we measured HR and BP variability. A decrease in HR variability (HRV) has been shown to predict cardiac mortality after myocardial infarction (48). HRV expressed as RMSSD obtained from telemeter-implanted, conscious mice showed no differences at 3 weeks after MI or any differences between genotypes (Fig. 5A). We also evaluated the contribution of different oscillatory components of HRV via power spectral analysis and found no differences in low frequency (Fig. 5B) or high frequency spectral powers (Fig. 5C). On the other hand, ASIC3^-/-^ mice had a significantly higher systolic blood pressure variability (BPV) than the WT mice after MI (Fig. 5D). We also found that the increase in LV mass from 48 hours to 3 weeks post-MI significantly correlated with BPV in ASIC3^-/-^ MI mice but not in WT MI mice (Fig. 5E).

**Figure 5.**
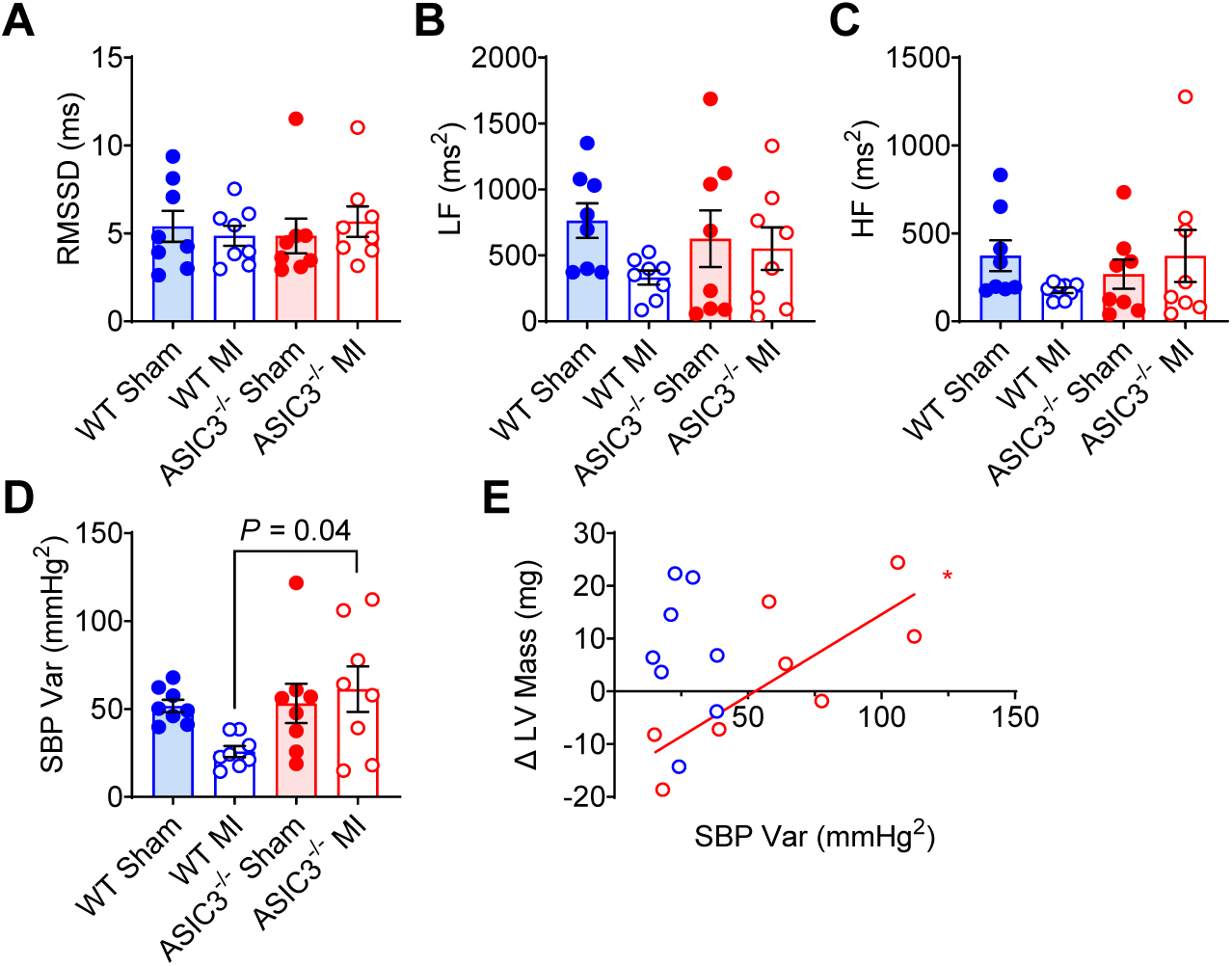
Heart rate and blood pressure variability. **A)** Root mean square of successive differences (RMSSD). Two-way ANOVA with Fisher’s LSD post-hoc test found no significant effects or individual differences. High frequency (HF) range **(B)** and low frequency (LF) range **(C)** from power spectra analysis. Two-way ANOVA with Fisher’s LSD post-hoc test found no significant effects or individual differences. **D)** Systolic blood pressure (SBP) variance. Two-way ANOVA revealed a significant effect of genotype (*F*(_1,28_) = 4.33, *P* = 0.047). Tukey post-hoc test found a significant difference between WT MI and ASIC3^-/-^ MI. **E)** Correlation between SBP variance and change in LV mass. Simple linear regression testing revealed a significant correlation for ASIC3^-/-^ MI (*R* = 0.79, *P* = 0.02, *) but not WT MI (*R* = 0.13, *P* = 0.8). (*N* = 8 for each group).

An increase in BPV is often associated with or driven by a decrease in baroreflex sensitivity. Baroreflex sensitivity can be used as a measure of autonomic responsiveness and is known to be diminished following myocardial infarction (49). Using our baseline telemetry data, we utilized spontaneous fluctuation in BP and HR to calculate the baroreflex gain (Fig. 6A) and engagement (Fig 6B) using the sequence technique in conscious, free-range mice. We did not see a diminishment in gain or engagement after MI in either group or any differences between genotypes. In addition to baroreceptor-heart rate reflex sensitivity, we also evaluated baroreceptor-renal sympathetic nerve activity (RSNA) reflex sensitivity in anesthetized mice by measuring the response of RSNA to changes in MAP evoked by infusion of varying doses of the vasodilator sodium nitroprusside followed by varying doses of the vasoconstrictor phenylephrine. ASIC3^-/-^ MI mice had reduced baroreceptor-RSNA reflex sensitivity compared to WT MI mice (Fig. 6C).

**Figure 6.**
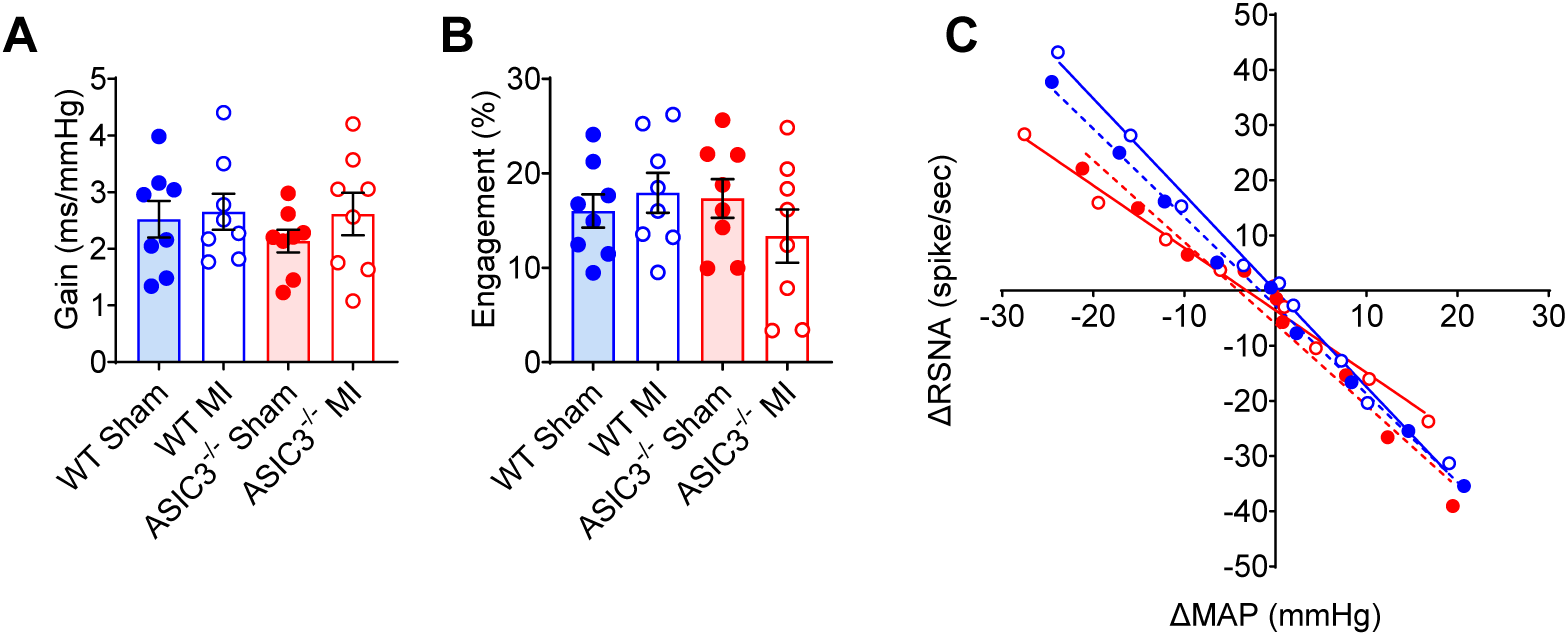
Blood pressure variability and baroreflex sensitivity. **A)** Baroreflex gain **(A)** and engagement **(B)** as measured via telemetry in conscious mice. Two-way ANOVA with Fisher’s LSD post-hoc test found no significant effects or individual differences. (*N* = 8 for each group). **C)** Baroreflex sensitivity determined by the response of renal sympathetic nerve activity (RSNA) to changes in mean arterial pressure (MAP) evoked by varying doses of sodium nitroprusside and phenylephrine. Two-way ANOVA of the slopes determined by simple linear regression revealed a significant effect of genotype (*F*(_1,325_) = 11.25, *P* = 0.0009) and a surgery x genotype interaction (*F*(_1,325_) = 5.05, *P* = 0.025). Tukey post-hoc test found significant differences between WT Sham and ASIC3^-/-^ MI (*P* = 0.023) and WT MI and ASIC3^-/-^ MI (*P* = 0.0003). (WT Sham: *N* = 8; WT MI: *N* = 14, ASIC3^-/-^ Sham: *N* = 9, ASIC3^-/-^ MI: *N* = 9).

To further elucidate genotype differences after MI, we performed qPCR on whole mouse hearts 3 weeks after surgery. As expected (50), there were no differences in the mRNA expression of alpha myosin heavy chain (*Myh6,* Fig 7A) and significant elevations in the beta myosin heavy chain (*Myh7,* Fig 7B) as well as the ratio of beta to alpha myosin heavy chains (*Myh7/Myh6,* Fig 7C) following MI in both genotypes, with no differences between the MI groups. There were also significant increases in both genotypes after MI in type 1 (*Col1a2,* Fig 7D) and type 3 (*Col3a1,* Fig 7E) collagen with no difference between the MI groups. While both MI groups had elevated expression levels of the brain natriuretic peptide (*Nppb,* Fig 7G), the ASIC3^-/-^ MI group had a marked increase in the expression of atrial natriuretic peptide (*Nppa*) compared to other groups (Fig. 7F).

**Figure 7.**
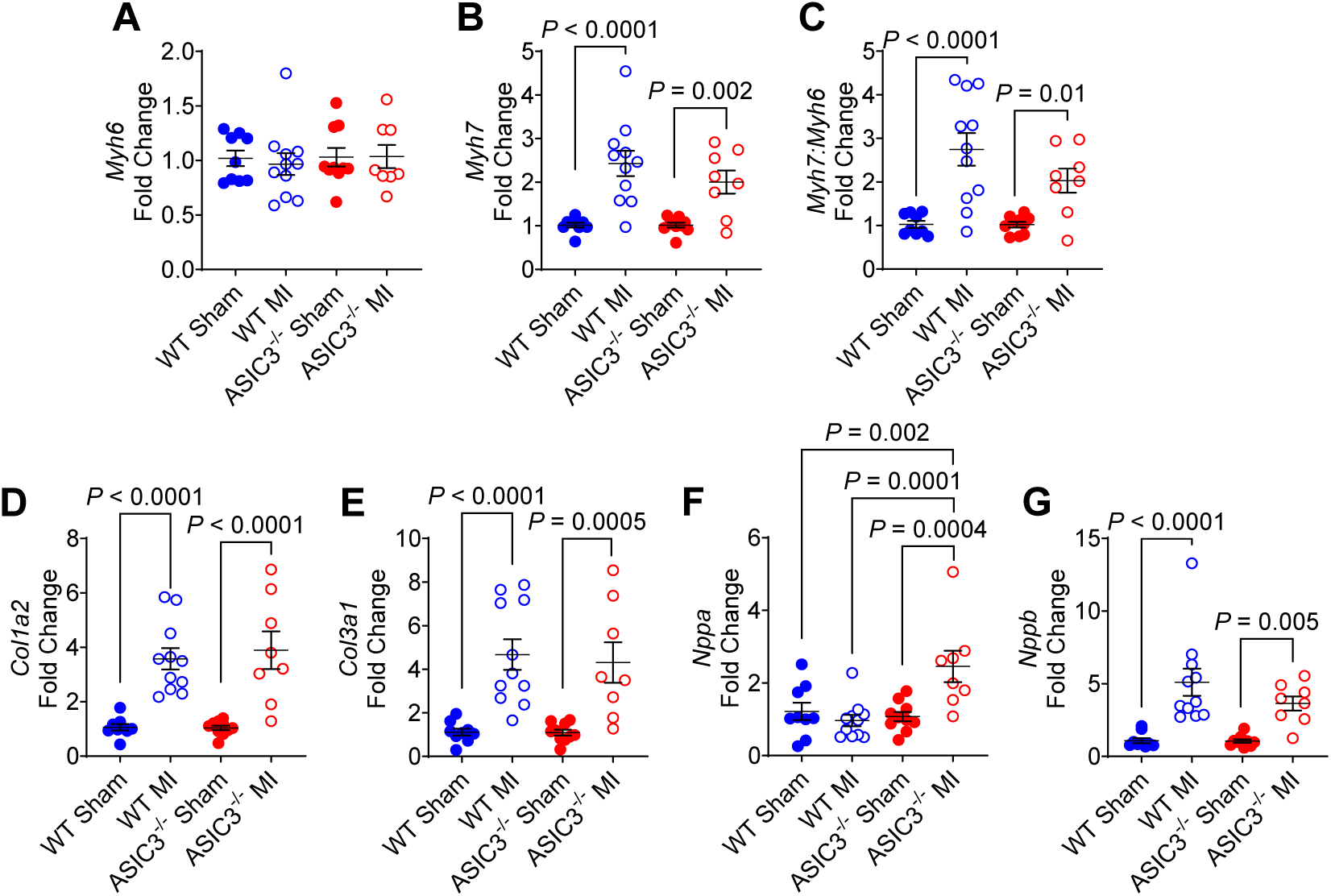
Quantitative PCR of whole mouse hearts. **A)** α-myosin heavy chain (*Myh6*). **B)** β-myosin heavy chain (*Myh7*). Two-way ANOVA with Fisher’s LSD post-hoc adjustment revealed a significant effect of surgery (F(_1,34_) = 33.71, *P* < 0.0001), with significant differences between WT Sham and WT MI, as well as ASIC3^-/-^ Sham and ASIC3^-/-^ MI. **C)** Ratio of β-myosin heavy chain to α-myosin heavy chain (*Myh7:Myh6*). Two-way ANOVA with Fisher’s LSD post-hoc adjustment revealed a significant effect of surgery (F(_1,34_) = 28.97, *P* < 0.0001), and significant differences between WT Sham and WT MI, and ASIC3^-/-^ Sham and ASIC3^-/-^ MI. **D)** Collagen type 1 (*Col1a2*). Two-way ANOVA with Fisher’s LSD post-hoc adjustment revealed a significant effect of surgery (F(_1,34_) = 50.8, *P* < 0.0001), with significant differences between WT Sham and WT MI, and ASIC3^-/-^ Sham and ASIC3^-/-^ MI. **E)** Collagen type 3 (*Col3a1*). Two-way ANOVA with Fisher’s LSD post-hoc adjustment revealed a significant effect of surgery (F(_1,34_) = 34.5, *P* < 0.0001), with significant differences between WT Sham and WT MI, and ASIC3^-/-^ Sham and ASIC3^-/-^ MI. **F)** Atrial natriuretic peptide (*Nppa*). Two-way ANOVA with Fisher’s LSD post-hoc adjustment revealed a significant effect of surgery (F(_1,34_) = 5.43, *P* = 0.03), genotype (F(_1,34_) = 7.73, *P* = 0.009), and their interaction (F(_1,34_) = 11.37, *P* = 0.002), with significant differences between ASIC3^-/-^ MI and all other groups. **G)** Brain natriuretic peptide (*Nppb*). Two-way ANOVA with Fisher’s LSD post-hoc adjustment revealed a significant effect of surgery (F(_1,34_) = 30.8, *P* < 0.0001), with differences between WT Sham and WT MI, and ASIC3^-/-^ Sham and ASIC3^-/-^ MI. Values are reported as mean ± SEM normalized to respective shams. (WT Sham: *N* = 9; WT MI: *N* = 11, ASIC3^-/-^ Sham: *N* = 10, ASIC3^-/-^ MI: *N* = 8).

## DISCUSSION

We tested the hypothesis that ASICs within peripheral sensory neurons may contribute to neurohormonal activation and adverse cardiac remodeling following MI. The major finding of this study is that following MI, ASIC3^-/-^ mice developed less LV dilation and increased LV hypertrophy, which may be driven by decreased baroreflex sensitivity and subsequent elevated BPV.

### Where and how is ASIC3 affecting cardiac remodeling after MI?

Since we studied a global knock-out model of ASIC3 deletion, an important follow-up question is where and how ASIC3 contributes to cardiac remodeling after MI. Whereas other ASIC subunits are expressed in both peripheral neurons and the central nervous system (CNS), ASIC3 is limited to peripheral sensory neurons in rodents (37, 51). Thus, the effects of ASIC3 deletion on cardiac remodeling likely involves the peripheral nervous system rather than direct CNS effects. We have shown that ASIC3 is expressed and probably plays a significant functional role in several afferent pathways, including cardiac and skeletal muscle afferents, as well as within the carotid body (28, 52, 53).

It is reasonable to believe that ASIC3 is sensing chemical/metabolic changes in and around the ischemic myocardium after infarction and, thereby, triggering reflexes that contribute to ventricular remodeling. During ischemia, the myocardium is exposed to rapid drops in pH. The interstitial pH, which is where cardiac afferent sensory nerve terminals lie, drops from 7.4 to ∼7.0 after coronary occlusion. Moreover, activation of cardiac sympathetic afferent fibers during ischemia correlates with the measured drop in pH, and addition of a pH buffer in the pericardial space significantly inhibits afferent activation (25). pH drops as small as 7.4 to 7.2 will cause sustained activation of ASIC3 currents within isolated cardiac afferents and generate action potentials (35, 54) – and as such, is particularly poised to sense these subtle pH changes within the heart during ischemia. Chronic dilated cardiomyopathy, even in the absence of significant epicardial coronary stenosis, is associated with regional and global myocardial ischemia due to microcirculatory abnormalities and increases in oxygen demand due to increases in wall stress (55, 56). We previously have shown that ASICs within cardiac sympathetic afferents are heteromeric channels composed of the ASIC3 and ASIC2a subunits. When ASIC3 is genetically deleted, the remaining ASIC2a forms homomeric channels that are insensitive to pH changes (34). Thus, loss of ASIC3 renders the cardiac afferent less capable of sensing metabolic changes within the heart.

Besides serving as local pH/metabolic sensors with cardiac afferents, ASICs might also serve as systemic pH sensors. As such, we found that ASICs function as pH sensors within glomus cells of the carotid body, which are the major peripheral chemoreceptors (53). Interestingly, the carotid body chemoreflex is augmented in a genetic rat model of hypertension, which correlated with an increase in ASIC3 expression and acid-evoked currents recorded from glomus cells (57). An exaggerated carotid body chemoreflex also contributes to sympathoexcitation in heart failure, and ablation of the carotid bodies in animal models of heart failure improves cardiac function and survival (15, 58).

ASICs are also expressed in arterial baroreceptors where they contribute to baroreflex sensitivity. We previously found that genetic deletion of ASIC2 in mice was associated with a diminishment of mechanical-induced depolarization in isolated cultured baroreceptor nodose neurons, a decrease in aortic depressor nerve activity to changes in arterial pressure in anesthetized mice, and diminished baroreflex gain and engagement in conscious mice (59). ASIC3 is also expressed in baroreceptors where it colocalizes with ASIC2 (59). Given the predilection of different ASIC subunits to heteromultimerize with each other (60), it is intriguing to speculate that ASIC3 subunits might also contribute to baroreceptive function. However, our results showed no difference in baroreceptor-heart rate and baroreceptor-RSNA reflex sensitivities between ASIC3^-/-^ and WT sham mice. In contrast, after MI, ASIC3^-/-^ mice had decreased baroreceptor-RSNA reflex sensitivity compared to WT mice (Fig. 6C). Various autonomic reflexes are intricately integrated and influence each other in multiple different ways (61–63). It is possible that ASICs are playing a role in cardiac remodeling through activation of several different sensory pathways, and future studies targeting ASICs within these specific sensory systems are likely to be insightful.

### Cardiac Remodeling

The degree of cardiac remodeling, and in particular LV dilation, is consistently one of the best predictors of poor clinical outcomes after MI (64–66). Therapeutics aimed to minimize and even reverse remodeling have shown to improve the symptoms of heart failure and improve survival (67). Both groups of animals, WT and ASIC3^-/-^, experienced remodeling after infarction. Here we show knocking out ASIC3 in mice reduced LV dilation that occurs in relation to the size of infarction and was associated with an increase in LV stroke volume. Additionally, ASIC3^-/-^ mice after MI had a significant increase in LV mass which, notably was significantly larger than that in WT mice. This suggests that WT mice post-MI developed eccentric hypertrophy – a relative increase in LV chamber size with less relative wall thickness. On the other hand, ASIC3^-/-^ mice developed concentric LV hypertrophy post-MI – a relative increase in wall thickness with less chamber enlargement. These differential effects of MI on remodeling in ASIC3^-/-^ compared to WT may explain the increase in LV stroke volume that was only observed in ASIC3^-/-^, but not in WT.

Concentric hypertrophy is often triggered by an increase in LV afterload, such as in the setting of hypertension or aortic stenosis. We found that MAP was similar between MI groups over a 48-hour period, however ASIC3^-/-^ mice had slightly higher BPs at night compared to WT mice after MI. This fits with our finding that ASIC3^-/-^ MI mice had an increase in BPV. We previously described that an increase in BPV generated in mice by sinoaortic baroreceptor denervation activates mechanosensitive pathways, including p125 focal adhesion kinase (p125-FAK) and p38 mitogen-activated protein kinase (p38-MAPK), resulting in cardiac hypertrophy, even though the average level of arterial BP was unchanged compared to sham-denervated mice (68). We postulated that higher BPV causes more periods of higher-than-average BP that triggers LV hypertrophy, and these periods of higher BP are not offset by periods of lower-than-normal BP. The baroreceptor denervation model of increased BPV was associated with increased collagen deposition within the LV at 12 weeks following baroreceptor denervation (68), whereas in our current study we did not find an increase in collagen mRNA at 3 weeks following MI. Perhaps the 3-week time point following MI was too early for a detectable increase in collagen mRNA expression to occur. Our results also contrast with other findings showing ASIC3 to be protective against fibrosis in a model of isoproterenol-induced myocardial ischemia (69). Another potential clue to explain the genotype differences in LV remodeling is that ASIC3^-/-^ MI mice had an increase in LV atrial natriuretic peptide that was not found in WT mice after MI. Local expression of atrial natriuretic peptide within the heart associates with LV hypertrophy (70).

### Autonomic nervous system changes in mice after MI

Myocardial infarction triggers a cascade of neurohormonal responses that resemble evolutionarily protected responses to hemorrhage or dehydration. In the latter cases the teleological “goal” is restoration of hemodynamic homeostasis. Along with an increase in sympathetic tone, MI can also lead to a decrease in parasympathetic tone. To our surprise, we found that WT mice did not develop the expected dysautonomia associated with MI. Injection of propranolol in conscious mice and SNA recordings in anesthetized mice did not demonstrate an increase in sympathetic tone following MI in WT mice. Similarly, we did not see a decrease in parasympathetic tone after atropine injection or in measurement of HRV. While mouse models have provided a robust platform to investigate cardiovascular disease, it is critical to understand the unique aspects of mouse physiology as they contrast to larger animal models and humans. In particular, the murine autonomic nervous system is characterized by a relatively high resting sympathetic tone and a lower parasympathetic tone, which has been shown to blunt the effects of sympathoexcitation after MI seen in other mammals (71–73). A logical next step of this work will be to modulate ASIC channels in an alternative animal model of myocardial infarction.

Previous work has shown the potential important contribution of cardiac afferents to disadvantageous cardiac remodeling after MI, and ablation of these sensory fibers using a toxin, resiniferatoxin, targeting TRPV1 channels improved remodeling after infarction (14, 21). While the clinical use of resiniferatoxin in a variety of clinical conditions is currently being investigated, the practicality of an invasive procedure involving the application of the toxin to the epicardial surface of the heart in patients suffering from MI is of question. A simpler strategy would be to find a molecular receptor within these sensory neurons that is activated after infarction. Interestingly, inhibiting TRPV1 does not seem to be the answer – genetic deletion of TRPV1 exacerbated maladaptive ventricular remodeling after MI (22). Here we demonstrate that genetic loss of ASIC3, a proton-activated ion channel principally expressed in sensory neurons, alters LV remodeling after MI in mice potentially by decreasing baroreflex sensitivity and increasing BPV. These results provide a strong rationale for follow-up studies investigating ASIC3 as a potential molecular target to treat post-MI ventricular remodeling.

## ACKNOWLEDGMENTS

We thank our lab member Tahsin Khataei and as well as Alan Ryan and Diane Olson for their input into the discussion of this work. We thank Mackenzie Spicer for her biostatistics expertise.

## NON-STANDARD ABBREVIATIONS AND ACRONYMNS

(MI): Myocardial infarction
(ASIC): Acid-sensing ion channel
(WT): Wild type
(BPV): Blood pressure variability
(sBPV): Systolic BPV
(SNA): Smypathetic nerve activity
(RSNA): Renal SNA
(spSNA): Splanchnic SNA
(RMSSD): Root mean square of successive differences
(LF): Low frequency
(HF): High frequency
(I.p.): Intraperitoneal
(IZ): Ischemic Zone
(IZ fx): IZ fraction
(BW): Body weight
(LVEDV): Left end diastolic volume
(LVESV): Left end systolic volume
(SV): Stroke volume
(EF): Ejection Fraction
(MAP): Mean arterial blood pressure
(BP): Blood pressure
(SBP): Systolic BPP
(DBP): Diastolic BP
(HR): Heart rate
(HRV): HR Variability
(BPV): BP variability

---

**Supplemental Table 1.** Cardiac remodeling as measured by the difference in echocardiographic data measured between 48 hours and 3 weeks, separated by sex.

|  |  | WT Sham | WT MI | ASIC3<sup>-/-</sup> Sham | ASIC3<sup>-/-</sup> MI |
| --- | --- | --- | --- | --- | --- |
| LVEDV (uL) | Male | 1.4 ± 4.2 | 29.2 ± 10.0 | -6.8 ± 4.2 | 22.3 ± 15.2 |
|  | Female | 1.6 ± 4.4 | 25.3 ± 9.0 | 2.3 ± 2.4 | 26.9 ± 6.5 |
| LVESV (uL) | Male | -0.3 ± 1.6 | 26.5 ± 9.0 | -1.2 ± 1.2 | 9.3 ± 12.9 |
|  | Female | 1.5 ± 1.8 | 22.3 ± 8.8 | 0.2 ± 0.4 | 18.5 ± 5.1 |
| LVEF | Male | 0.02 ± 0.03 | -0.04 ± 0.03 | -0.02 ± 0.02 | 0.13 ± 0.05 |
|  | Female | -0.02 ± 0.02 | -0.01 ± 0.05 | 0.00 ± 0.02 | 0.01 ± 0.03 |
| LV Mass<br>(mg) | Male | 0.8 ± 4.2 | 0.04 ± 4.2 | -1.1 ± 3.5 | 18.3 ± 5.4 |
|  | Female | -2.3 ± 2.8 | 6.2 ± 3.4 | 9.6 ± 4.8 | 9.8 ± 3.4 |
| SV (uL) | Male | 1.6 ± 3.4 | 2.8 ± 2.2 | -5.6 ± 3.4 | 12.9 ± 5.7 |
|  | Female | 0.06 ± 2.7 | 3.0 ± 3.2 | 2.1 ± 2.2 | 8.4 ± 2.9 |

Values are means ± SE; Statistical analysis by three-way ANOVA with Tukey post hoc correction revealed no statistical differences between the sexes for any of the following parameters: left ventricular end diastolic volume (LVEDV): $F_{(1,111)} = 0.1419$ , $P = 0.7071$ ; left ventricular end systolic volume (LVESV): $F_{(1,111)} = 0.1280$ , $P = 0.7211$ ; left ventricular ejection fraction (LVEF): $F_{(1,111)} = 0.518$ , $P = 0.2205$ ; left ventricular (LV) mass: $F_{(1,111)} = 0.1742$ , $P = 0.6772$ ; stroke volume (SV): $F_{(1,111)} = 0.03767$ , $P = 0.8465$ ; WT Sham male: $N = 10$ ; WT Sham female: $N = 11$ ; WT MI male: $N = 25$ ; WT MI female: $N = 14$ ; ASIC3<sup>-/-</sup> Sham male: $N = 12$ ; ASIC3<sup>-/-</sup> Sham female: $N = 12$ ; ASIC3<sup>-/-</sup> MI male: $N = 9$ ; ASIC3<sup>-/-</sup> MI female: $N = 26$ .

**Supplemental Table 2.**
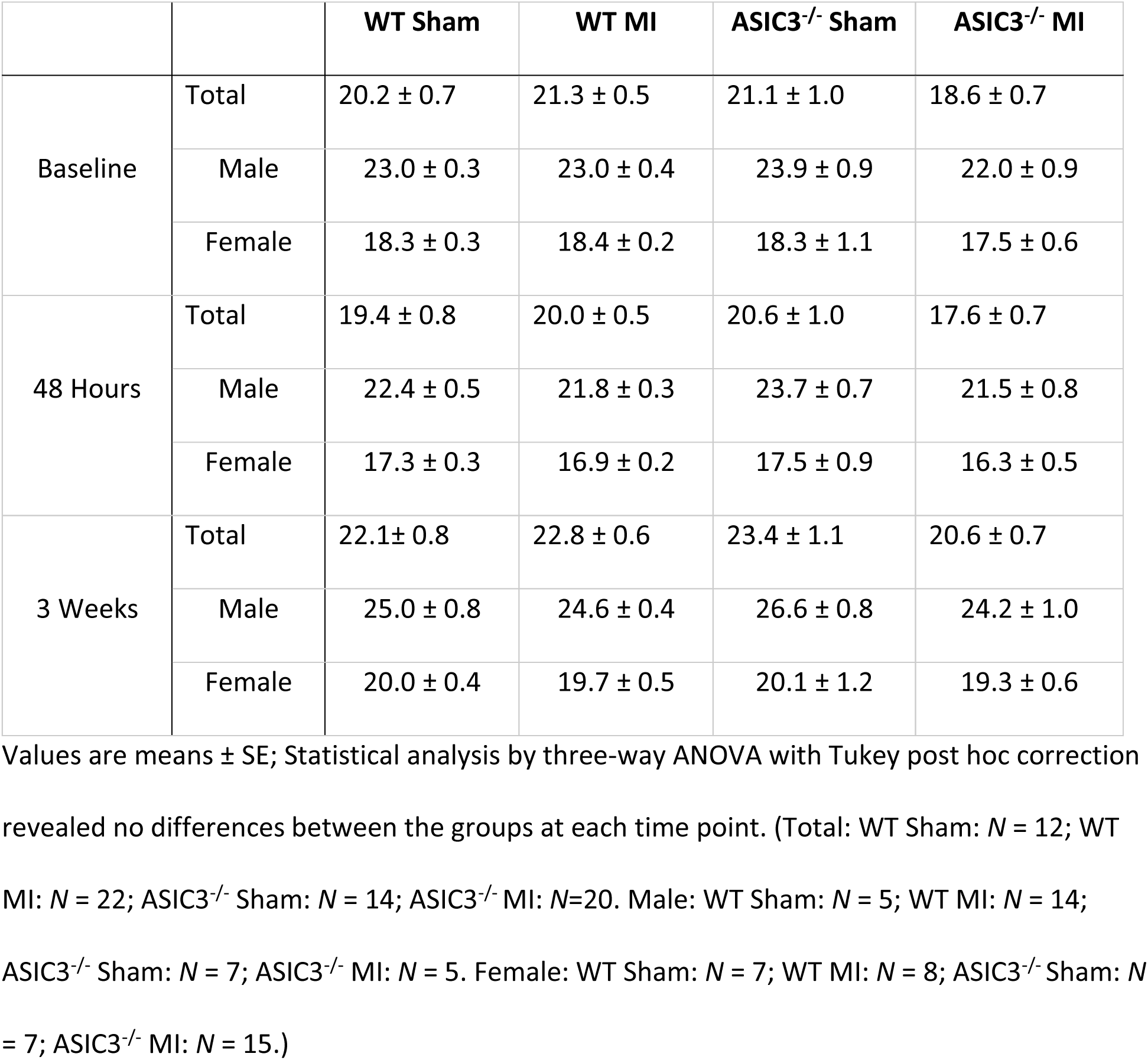
Body Weights (mg) measure at 3 weeks after myocardial infarction (MI) or sham surgery.

|  |  | **WT Sham** | **WT MI** | **ASIC3<sup>-/-</sup> Sham** | **ASIC3<sup>-/-</sup> MI** |
| --- | --- | --- | --- | --- | --- |
| Baseline | Total | 20.2 ± 0.7 | 21.3 ± 0.5 | 21.1 ± 1.0 | 18.6 ± 0.7 |
|  | Male | 23.0 ± 0.3 | 23.0 ± 0.4 | 23.9 ± 0.9 | 22.0 ± 0.9 |
|  | Female | 18.3 ± 0.3 | 18.4 ± 0.2 | 18.3 ± 1.1 | 17.5 ± 0.6 |
| 48 Hours | Total | 19.4 ± 0.8 | 20.0 ± 0.5 | 20.6 ± 1.0 | 17.6 ± 0.7 |
|  | Male | 22.4 ± 0.5 | 21.8 ± 0.3 | 23.7 ± 0.7 | 21.5 ± 0.8 |
|  | Female | 17.3 ± 0.3 | 16.9 ± 0.2 | 17.5 ± 0.9 | 16.3 ± 0.5 |
| 3 Weeks | Total | 22.1 ± 0.8 | 22.8 ± 0.6 | 23.4 ± 1.1 | 20.6 ± 0.7 |
|  | Male | 25.0 ± 0.8 | 24.6 ± 0.4 | 26.6 ± 0.8 | 24.2 ± 1.0 |
|  | Female | 20.0 ± 0.4 | 19.7 ± 0.5 | 20.1 ± 1.2 | 19.3 ± 0.6 |

Values are means ± SE; Statistical analysis by three-way ANOVA with Tukey post hoc correction revealed no differences between the groups at each time point. (Total: WT Sham: *N* = 12; WT MI: *N* = 22; ASIC3<sup>-/-</sup> Sham: *N* = 14; ASIC3<sup>-/-</sup> MI: *N* = 20. Male: WT Sham: *N* = 5; WT MI: *N* = 14; ASIC3<sup>-/-</sup> Sham: *N* = 7; ASIC3<sup>-/-</sup> MI: *N* = 5. Female: WT Sham: *N* = 7; WT MI: *N* = 8; ASIC3<sup>-/-</sup> Sham: *N* = 7; ASIC3<sup>-/-</sup> MI: *N* = 15.)

**Supplemental Table 3.**
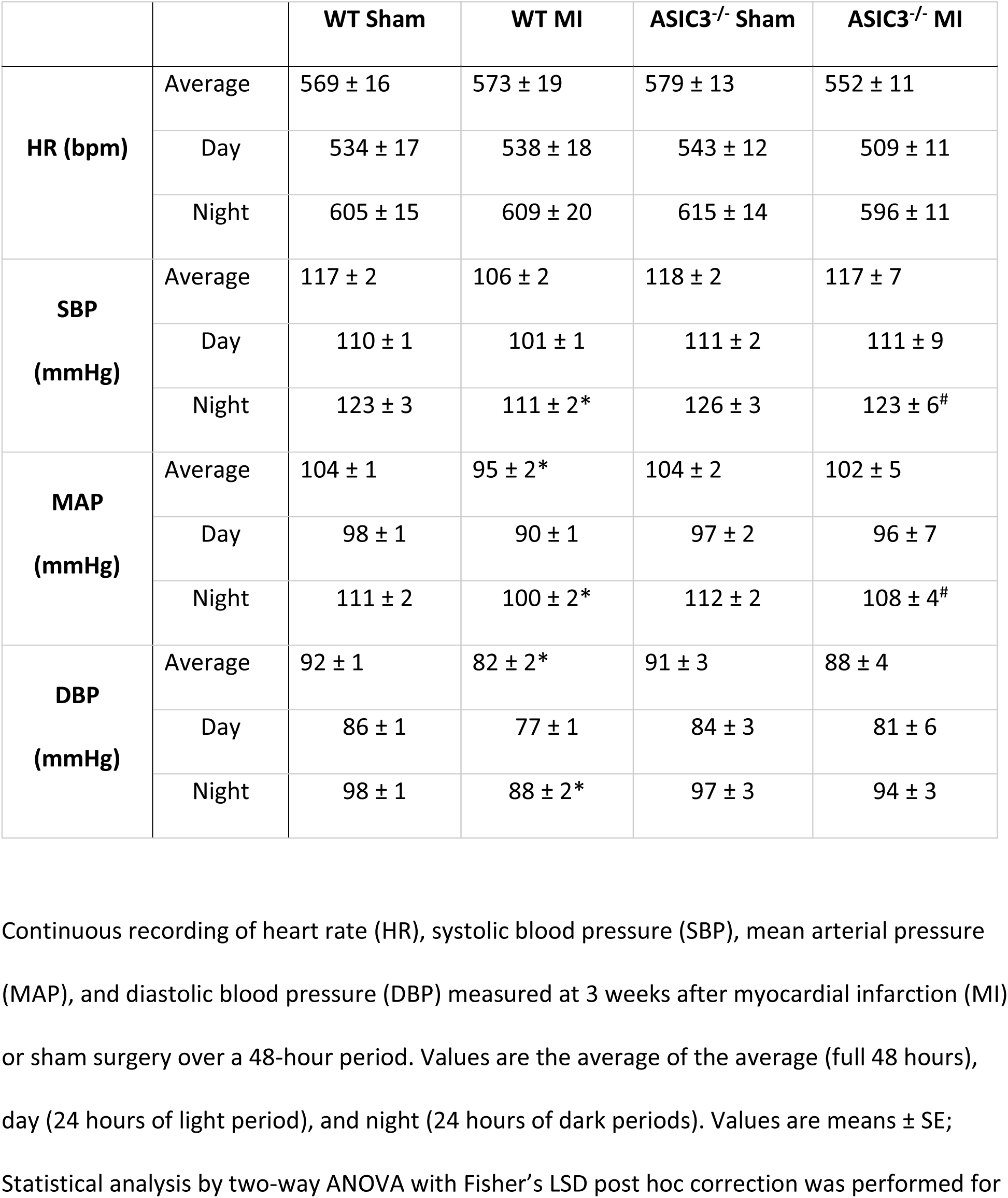
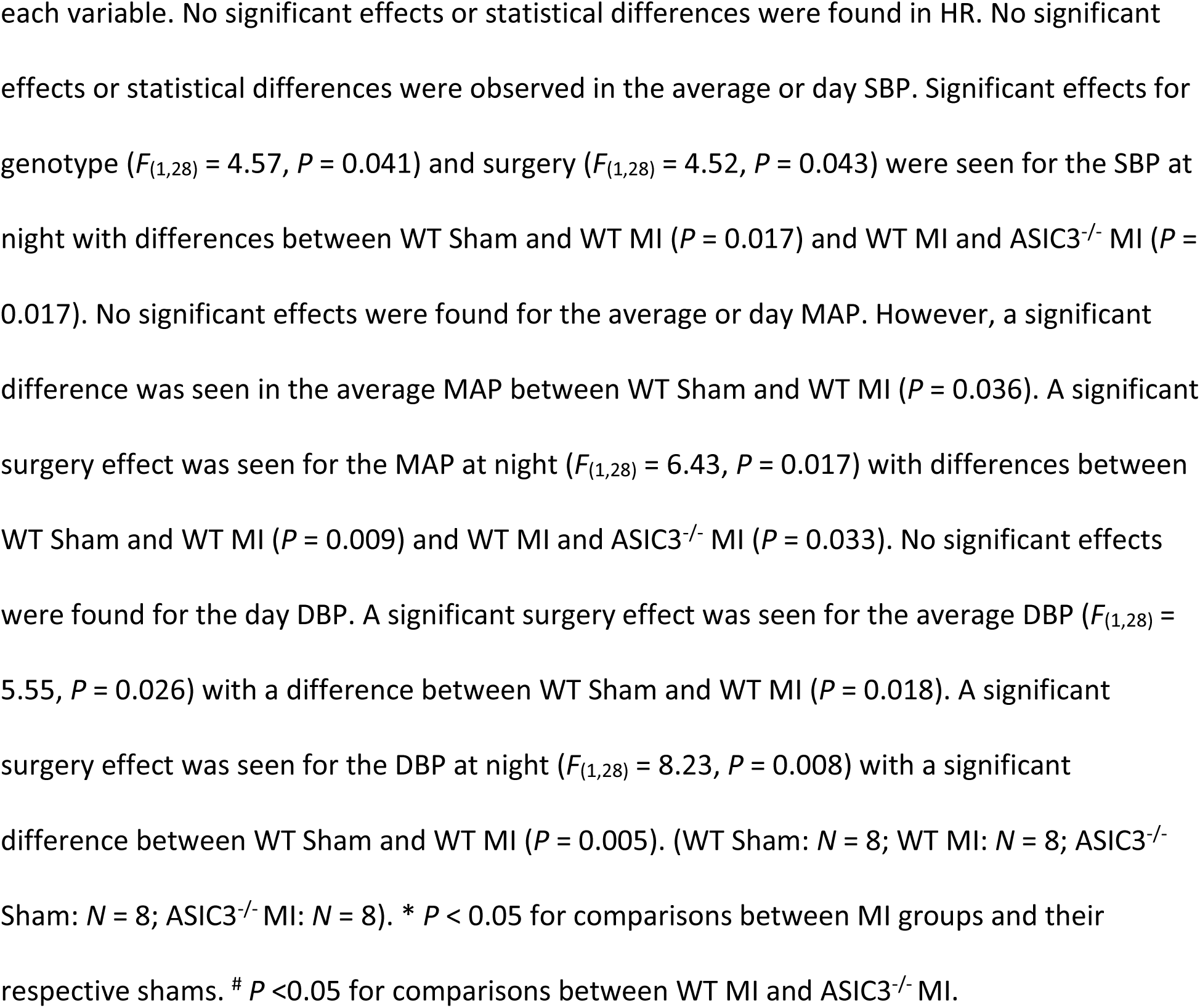
Hemodynamics measured over 48 hours at 3 weeks after myocardial infarction (MI) or sham surgery.

|  |  | **WT Sham** | **WT MI** | **ASIC3<sup>-/-</sup> Sham** | **ASIC3<sup>-/-</sup> MI** |
| --- | --- | --- | --- | --- | --- |
| **HR (bpm)** | Average | 569 ± 16 | 573 ± 19 | 579 ± 13 | 552 ± 11 |
|  | Day | 534 ± 17 | 538 ± 18 | 543 ± 12 | 509 ± 11 |
|  | Night | 605 ± 15 | 609 ± 20 | 615 ± 14 | 596 ± 11 |
| **SBP (mmHg)** | Average | 117 ± 2 | 106 ± 2 | 118 ± 2 | 117 ± 7 |
|  | Day | 110 ± 1 | 101 ± 1 | 111 ± 2 | 111 ± 9 |
|  | Night | 123 ± 3 | 111 ± 2* | 126 ± 3 | 123 ± 6<sup>#</sup> |
| **MAP (mmHg)** | Average | 104 ± 1 | 95 ± 2* | 104 ± 2 | 102 ± 5 |
|  | Day | 98 ± 1 | 90 ± 1 | 97 ± 2 | 96 ± 7 |
|  | Night | 111 ± 2 | 100 ± 2* | 112 ± 2 | 108 ± 4<sup>#</sup> |
| **DBP (mmHg)** | Average | 92 ± 1 | 82 ± 2* | 91 ± 3 | 88 ± 4 |
|  | Day | 86 ± 1 | 77 ± 1 | 84 ± 3 | 81 ± 6 |
|  | Night | 98 ± 1 | 88 ± 2* | 97 ± 3 | 94 ± 3 |

Continuous recording of heart rate (HR), systolic blood pressure (SBP), mean arterial pressure (MAP), and diastolic blood pressure (DBP) measured at 3 weeks after myocardial infarction (MI) or sham surgery over a 48-hour period. Values are the average of the average (full 48 hours), day (24 hours of light period), and night (24 hours of dark periods). Values are means ± SE; Statistical analysis by two-way ANOVA with Fisher's LSD post hoc correction was performed for
each variable. No significant effects or statistical differences were found in HR. No significant effects or statistical differences were observed in the average or day SBP. Significant effects for genotype ( $F_{(1,28)} = 4.57, P = 0.041$ ) and surgery ( $F_{(1,28)} = 4.52, P = 0.043$ ) were seen for the SBP at night with differences between WT Sham and WT MI ( $P = 0.017$ ) and WT MI and ASIC3<sup>-/-</sup> MI ( $P = 0.017$ ). No significant effects were found for the average or day MAP. However, a significant difference was seen in the average MAP between WT Sham and WT MI ( $P = 0.036$ ). A significant surgery effect was seen for the MAP at night ( $F_{(1,28)} = 6.43, P = 0.017$ ) with differences between WT Sham and WT MI ( $P = 0.009$ ) and WT MI and ASIC3<sup>-/-</sup> MI ( $P = 0.033$ ). No significant effects were found for the day DBP. A significant surgery effect was seen for the average DBP ( $F_{(1,28)} = 5.55, P = 0.026$ ) with a difference between WT Sham and WT MI ( $P = 0.018$ ). A significant surgery effect was seen for the DBP at night ( $F_{(1,28)} = 8.23, P = 0.008$ ) with a significant difference between WT Sham and WT MI ( $P = 0.005$ ). (WT Sham: $N = 8$ ; WT MI: $N = 8$ ; ASIC3<sup>-/-</sup> Sham: $N = 8$ ; ASIC3<sup>-/-</sup> MI: $N = 8$ ). \* $P < 0.05$ for comparisons between MI groups and their respective shams. # $P < 0.05$ for comparisons between WT MI and ASIC3<sup>-/-</sup> MI.

